# Dietary fiber content in clinical ketogenic diets modifies the gut microbiome and seizure resistance in mice

**DOI:** 10.1101/2024.07.31.606041

**Authors:** Ezgi Özcan, Kristie B. Yu, Lyna Dinh, Gregory R. Lum, Katie Lau, Jessie Hsu, Mariana Arino, Jorge Paramo, Arlene Lopez-Romero, Elaine Y. Hsiao

## Abstract

The gut microbiome is emerging as an important modulator of the anti-seizure effects of the classic ketogenic diet. However, many variations of the ketogenic diet are used clinically to treat refractory epilepsy, and how different dietary formulations differentially modify the gut microbiome in ways that impact seizure outcome is poorly understood. We find that clinically prescribed ketogenic infant formulas vary in macronutrient ratio, fat source, and fiber content and also in their ability to promote resistance to 6-Hz psychomotor seizures in mice. By screening specific dietary variables for their effects on a model human infant microbial community, we observe that dietary fiber, rather than fat ratio or source, drives substantial metagenomic shifts. Addition of dietary fiber to a fiber-deficient ketogenic formula restores seizure resistance, and supplementing protective ketogenic formulas with excess dietary fiber further potentiates seizure resistance. By screening 13 fiber sources and types, we identify distinct subsets of metagenomic responses in the model human infant microbial community that correspond with increased seizure resistance in mice. In particular, supplementation with seizure-protective fibers enriches microbial representation of genes related to queuosine biosynthesis and preQ_0_ biosynthesis and decreases representation of microbial genes related to sucrose degradation, which is also seen in seizure-protected mice that are fed fiber-containing ketogenic infant formulas. Overall, this study reveals that different formulations of clinical ketogenic diets, and dietary fiber content in particular, differentially impact seizure outcome in mice, likely through modification of the gut microbiome. Understanding interactions between dietary components of the ketogenic diet, the gut microbiome, and host susceptibility to seizures could inform novel microbiome-guided approaches to treat refractory epilepsy.

## Introduction

The low-carbohydrate, high-fat ketogenic diet (KD) is used to treat epilepsy in children who do not respond positively to existing anti-seizure medications. While it is well integrated into the healthcare system, KD therapies have variable effectiveness in reducing seizures, ranging from 45 to 85% in infants and children that exhibit high compliance^1–5^ and with substantially lower rates in adults^6,7^. Recent reports highlight a key role for the gut microbiome in mediating effects of the KD on various host physiologies, including glucose and lipid metabolism^8^, immune function^9,10^, brain activity^11^, and behavior^10,12,13^. The KD alters the gut microbiome across several human and animal epilepsy studies^14–19^, and relationships are seen between the gut microbiome and seizure resistance in various rodent epilepsy models^12,20–23^. Findings from the field are converging upon the notion that variation in the gut microbiome may contribute to variability in patient responsiveness to the KD, and that microbiome-targeted interventions could be used to promote the efficacy of the KD in treating refractory epilepsy.

While existing studies of the microbiome and KD have focused predominantly on the classic KD, many variations of the KD with different macronutrient ratios and types are used clinically to treat epilepsy, depending on factors such as the age of the patient, seizure type, and tolerability of the dietary regimen^24–27^. For example, the KD is commonly administered as a 4:1 or 3:1 fat to carbohydrate and protein ratio, depending on patient tolerance. The medium chain triglyceride (MCT) diet, often derived from MCT-rich coconut oil, is thought to promote enhanced ketone production while being less restrictive than the classic KD. The Modified Atkins Diet (MAD), which does not require strict weighing of food or fluids, and Low Glycemic Index Treatment (LGIT), which focuses on carbohydrates with low glycemic index rather than removal, are additional less restrictive variations of the KD that are frequently used in older children and adults.

While only a few small human studies have compared different KD variants for their seizure reduction, citing no significant differences^25,26,28,29^, other research suggests that differences in dietary formulation may impact host responses to the KD. In a retrospective open label trial of patients with drug resistant epilepsy, transitioning to a polyunsaturated fatty acid (PUFA)-based KD enhanced seizure control in individuals who responded poorly to the classic KD^30^. Moreover, differences in dietary formulation can have substantial impacts on microbiome-dependent host phenotypes – KDs with different fat ratio and/or source resulted in differential influences of the microbiome on host glucose and lipid metabolism, as well as immune function^8,9^. In addition, supplementation with dietary fiber, a key energy source for gut bacteria that modulates myriad host metabolic, immune, and neural functions, is incorporated into some clinical KD regimens to ease gastrointestinal symptoms^31^, but whether it alters seizure response is unclear^32^. Overall, increasing research indicating that the gut microbiome modifies seizure susceptibility and the anti-seizure effects of the KD raises the important question of how variations in the formulation of medical KDs differentially shape the microbiome in ways that impact seizure outcome.

In this study, we tested effects of three clinically prescribed KD infant formulas on the mouse gut microbiome and resistance to 6-Hz psychomotor seizures, as a benchmark model of refractory epilepsy^33^. To determine which dietary variables serve as key drivers of microbiome response, we established a model human infant microbial consortium and assessed effects of fat ratio, fat source, and carbohydrate source on shaping its functional potential. We further screened 13 fiber sources and types for their differential impacts on the model infant microbial community and tested top candidates for their ability to restore and/or potentiate seizure protective effects of clinical KD infant formulas. Results from this study reveal key diet-microbiome interactions that promote the seizure protective effects of medical KDs.

## Results

### Different clinical KD infant formulas elicit differential seizure responses in mice

Mechanistic studies of the KD on seizure resistance often rely on commercial KD chows that are formulated for lab animals and not directly relevant to medical KD therapies used for human epilepsy. At the same time, clinical KD regimens vary widely in nutritional content and are often tailored to the particular individual’s needs and tolerability, making it difficult to identify standard regimens. To examine how clinically relevant formulations of the KD elicit differential effects on seizure outcome, we focused on 3 commonly prescribed commercial KD infant formulas – KD4:1, KD3:1, and MCT2.5:1 -- due to their reproducible composition, direct clinical relevance, frequent prescription, and importance for infants and young children as especially vulnerable subsets of refractory epilepsy patients for which improved interventions are needed. Compared to a standard infant formula as a control diet (CD), the 3 KD infant formulas all exhibit high fat content relative to carbohydrate and protein, but they display nuanced differences in formulation (**Fig. 1a, Supplementary Data 1**). In addition to differences in fat ratio, fat source varies between the formulations, where KD4:1 contains soy lecithin but lacks coconut oil (MCT source) and linoleic acid, KD3:1 contains linoleic acid but lacks soy lecithin and coconut oil, and MCT2.5:1 contains coconut oil but lacks soy lecithin and linoleic acid. There are also differences in carbohydrate content, where both KD4:1 and MCT2.5:1 contain corn syrup solids, high amylose corn starch, chicory root inulin, gum arabic, cellulose, fructooligosaccharides (FOS), soy fiber, and maltodextrin, whereas KD3:1 contains only lactose and corn syrup solids, with none of the dietary fibers. The CD contains lactose and less than 2% dietary fiber comprised of galactooligosaccharides, which differs from the types of fibers included in KD4:1 and MCT2.5:1.

**Fig 1.**
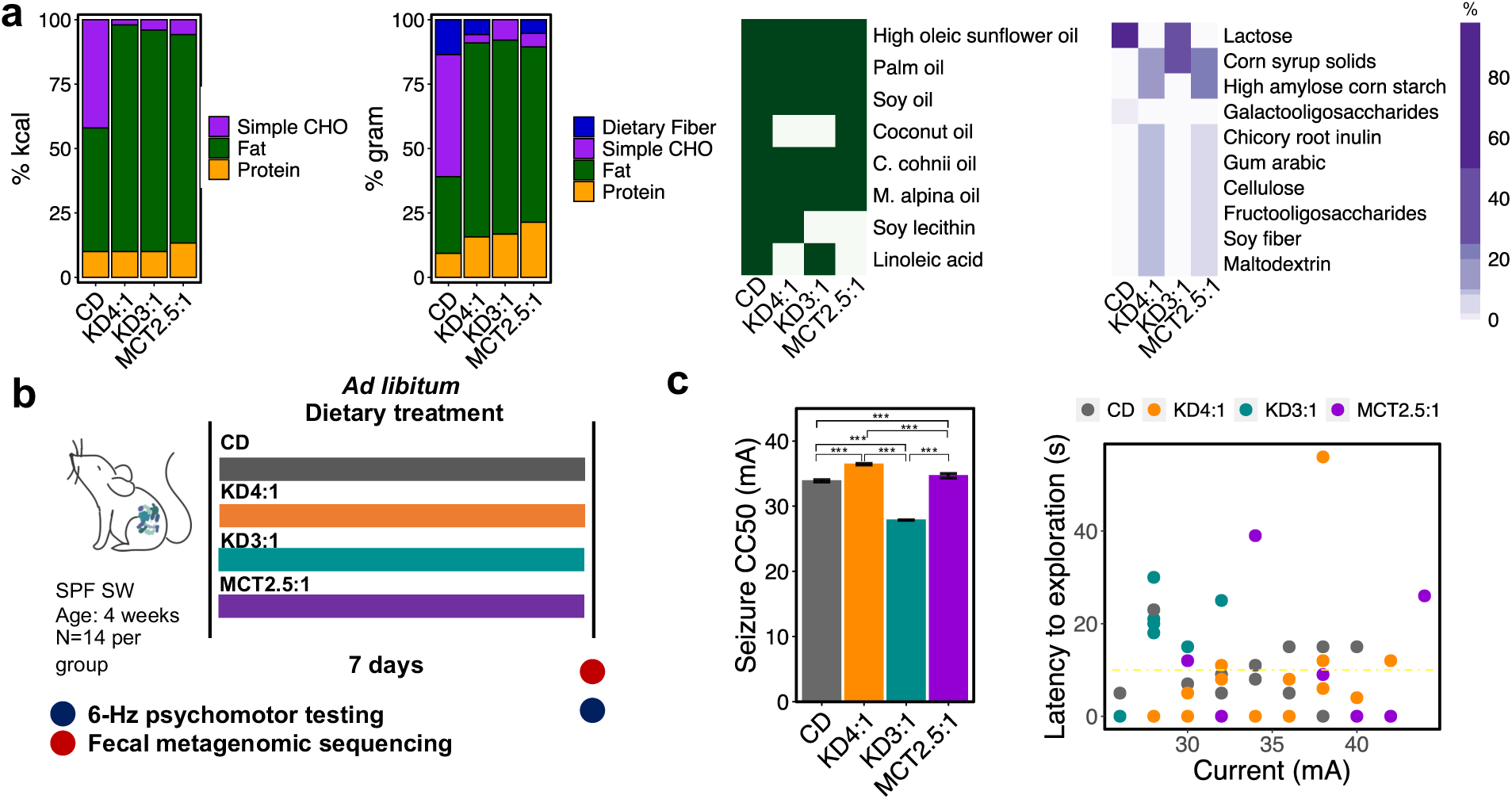
Different formulations of medical ketogenic diets (KD) elicit differential responses to 6-Hz seizures in mice. a. Macronutrient composition without fiber (for determining KD fat ratio), macronutrient composition with fiber, absence/presence of fat sources, and percent carbohydrate composition for the commercial KD infant formulas KD4:1, KD3:1, and MCT2.5:1, relative to standard infant formula as control diet (CD). b. Experimental design: 4 week old conventional (specific pathogen free, SPF) Swiss Webster (SW) mice (n=14 mice/group) were fed each medical KD or CD as liquid diets for 7 days. c. 6-Hz seizure threshold (left) and latency to exploration (right) for mice fed KDs or CD as liquid diet (left, one-way ANOVA with Bonferroni, n=14 mice/group, ***p<0.001). Yellow line at y = 10 s represents threshold for scoring seizures.

To determine how different KD formulations impact seizure susceptibility, we fed cohorts of conventional 4 week-old mice the KD4:1, KD3:1, MCT2.5:1, or CD formula as liquid diet for 1 week, and then tested for susceptibility to 6-Hz psychomotor seizures (**Fig. 1b**). Juvenile mice were selected to mimic the typical use of the KD to treat pediatric epilepsy, to align the timing of mouse brain development to early brain development in humans^34^, and to preclude effects of pre-weaning treatment, where effects of the diets on maternal behavior and physiology would confound their direct effects on offspring. 1 week of feeding was selected based on our prior longitudinal characterization, which indicated that KD chow shifts the gut microbiome and confers seizure protection by day 4 of treatment in mice^12^. Finally, the 6-Hz seizure assay was selected as a benchmark model of refractory epilepsy that is used to screen for new anti-seizure medications^33^ and involves low-frequency corneal stimulation to induce complex partial seizures related to human temporal lobe epilepsy^35^. KD chow protects against 6-Hz seizures, as indicated by increases in current intensity required to elicit a seizure in 50% of the subjects tested (CC50, seizure threshold)^12,36,37^.

As seen previously for KD chows^12,36,37^, we observed that feeding mice clinical KD4:1 infant formula increased seizure thresholds compared to controls fed a CD infant formula (**Fig. 1c**). MCT2.5:1 also increased seizure thresholds albeit to a lesser degree than KD4:1, which may be due to its comparatively lower fat ratio or different fat source. In contrast, however, KD3:1 infant formula yielded decreased seizure thresholds compared to all other groups, including CD-fed controls, suggesting that the KD3:1 formulation increases susceptibility to 6-Hz seizures in mice. There was no correlation of seizure threshold with average calories consumed for the different KDs or with degree of ketosis as assessed by serum levels of beta-hydroxybutyrate (**Supplementary** Fig. 1a-b). To further assess whether the differences in seizure outcome may be confounded by nuances of providing the diet in liquid form, such as differences in density or leakage from the bottle, we repeated the experiment by providing the infant formula diets in solid form following dehydration. Consistent with our previous observation, solid KD4:1 and MCT2.5:1 increased seizure threshold relative to controls fed solid CD, whereas solid KD3:1 decreased resistance to 6-Hz seizures, with no correlation with total diet consumed (**Supplementary** Fig. 1c, d). These data indicate that variations in clinical KD formulations differentially modify host resistance versus susceptibility to 6-Hz seizures in mice.

### Clinical KD infant formulas differentially alter the mouse gut microbiome

Classic KD-induced changes in the mouse and human microbiome are necessary and/or sufficient to confer resistance to 6-Hz seizures in mice^12,20^. To determine how the different clinical KD infant formulas impact the gut microbiome, we performed metagenomic sequencing of fecal microbiota from mice fed KD4:1, KD3:1, MCT2.5:1, or CD for 1 week. In contrast to results from KD vs. standard chow^12,38^, KD4:1 and MCT2.5:1 significantly increased α-diversity of the microbiome, as indicated by elevated Shannon’s diversity index, when compared to CD controls (**Fig. 2a**). However, there was no significant effect of KD3:1 on Shannon diversity levels, despite comparable increases across all KD formula groups in species richness of the fecal microbiota. This suggests that the main driver of α-diversity differences between the KD groups is differential alteration in species evenness—indeed, KD3:1 yielded fecal microbiota with significantly reduced Pielou’s evenness compared to KD4:1 and MCT2.5:1 groups. β-diversity analysis of the gut microbiota based on Bray-Curtis dissimilarity and weighted Unifrac distances showed that KD samples clustered distinctly from CD controls along PCoA1, with KD4:1 and MCT2.5:1 samples showing further separation from CD than KD3:1 samples (PERMANOVA, p=0.001, R^2^=0.6, **Fig. 2b**). In particular, all KD groups exhibited significantly decreased relative abundances of *Actinobacteria* and increased *Bacteroidetes* and unclassified bacteria compared to CD controls (**Supplementary** Fig. 2a). However, only KD4:1 and MCT2.5:1 shared statistically significant decreases in *Erysipelotrichia* and increases in *Streptococcaceae*, *Coriobacteriia*, and *Deferribacteres*, whereas KD3:1 exhibited no significant changes in these taxa compared to CD (**Supplementary** Fig. 2a-c). Rather, KD3:1 showed significantly increased relative abundance of *Proteobacteria*, *Escherichia coli, Enterococcus faecalis,* and *Mammaliicoccus sciuri* compared to CD, KD4:1, and/or MCT2.5:1 (**Supplementary** Fig. 2a-c).

**Figure 2:**
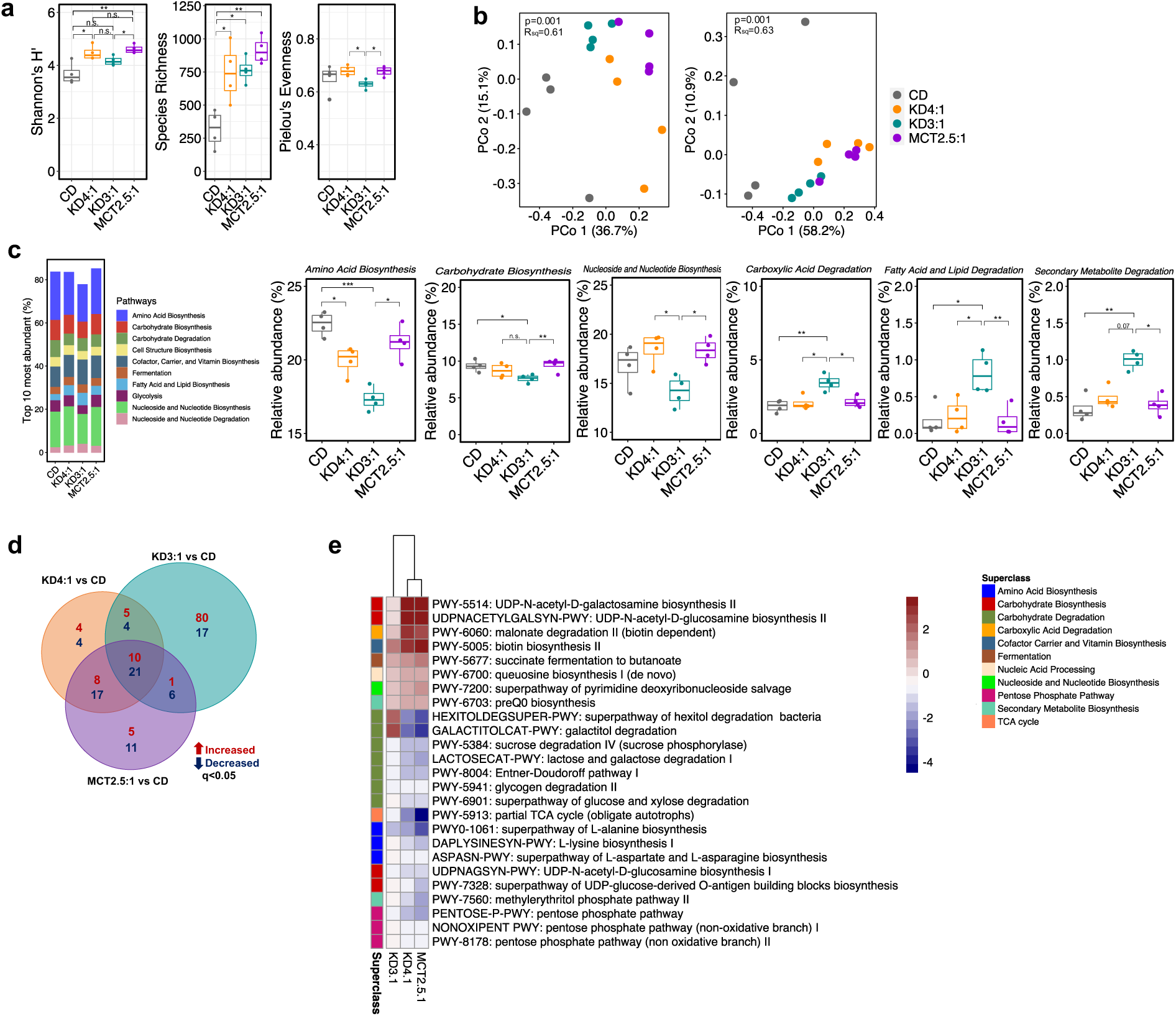
Medical KDs induce differential alterations in the gut microbiome that associate with resistance vs. susceptibility to 6-Hz seizures. a. Alpha diversity from fecal metagenomic sequencing data after treatment with KDs or CD (Kruskal-Wallis with Dunn’s test: ^∗^p < 0.05,**p <0.01 n.s., not statistically significant; n=4 cages/group. Data are presented as box-and-whisker plots with median and first and third quartiles). b. Principal coordinates analysis (PCoA) of Bray-Curtis dissimilarity (left) and weighted UniFrac distance (right) based on fecal metagenomic sequencing data after dietary treatment. (PERMANOVA, n = 4 cages/group). c. Top 10 most abundant metagenomic superclass pathways (left). Differentially abundant pathways that are significantly altered in seizure susceptible group KD3:1 and/or shared between seizure protected groups KD4:1 and MCT2.5:1. (Kruskal-Wallis with Dunn’s test: ^∗^p < 0.05, **p< 0.01, ***p<0.001; n=4 cages/group. Data are presented as box-and-whisker plots with median and first and third quartiles). d. Venn diagram of differential metagenomic pathways (q<0.05) for each KD relative to CD. (MaAsLin2, General Linear Model (GLM); n=4 cages/group). e. Heatmap of differential metagenomic pathways (q<0.05) that are shared between seizure-protected groups KD4:1 and MCT2.5:1 and not significant in seizure-susceptible group KD3:1. (GLM statistical test, n=4/condition)

The seizure susceptible KD3:1 group also exhibited decreased representation of the top 10 most abundant metagenomic superclass pathways (**Fig. 2c**), suggesting that the KD3:1 limits the presence of microbial taxa associated with prevalent functions and/or enriches the representation of previously rare metagenomic pathways. Among the top 10, the relative abundance of superclass pathways related to amino acid, carbohydrate, and nucleoside and nucleotide biosyntheses were significantly lower in KD3:1 relative to MCT2.5:1, CD, and/or KD4:1 groups. In contrast, superclass pathways related to carboxylic acid, fatty acid and lipid, and secondary metabolite degradation were significantly elevated in KD3:1 compared to other groups. When considering specific alterations at the more resolved pathway level, all three KDs shared subsets of metagenomic changes compared to CD controls, where KD4:1 and MCT2.5:1 shared greater overlap than with KD3.1 (**Fig 2d**). Namely, KD4:1 and MCT2.5:1 (but not KD3:1) similarly induced significant metagenomic increases in select pathways related to carbohydrate biosynthesis (UDP-N-acetyl-D-galactosamine II and UDP-N-acetyl-D-glucosamine biosynthesis II), carboxylic acid degradation (biotin-dependent malonate degradation), and cofactor, carrier, and vitamin biosynthesis (biotin biosynthesis), and decreases in select pathways related to carbohydrate degradation (hexitol and galactitol degradation, sucrose, lactose, galactose degradation, and Entner-Doudoroff pathway), amino acid biosynthesis (L-lysine and L-alanine biosynthesis), carbohydrate biosynthesis (UDP-N-acetyl-D-glucosamine biosynthesis I and UDP-glucose-derived-O-antigen building blocks biosynthesis), and pentose phosphate pathway compared to CD controls (**Fig. 2e**). KD3:1 displayed the most differentially abundant metagenomic pathways compared to CD, which were distinct from those seen in the other KD groups (Supplementary Fig. 2d). The majority of differentially abundant pathways that were elevated by KD3:1 related to amide, amidine, amine, and polyamine degradation, fatty acid and lipid biosynthesis, carboxylic acid degradation, and fermentation (Supplementary Fig. 2d). In particular, pathways for phospholipid remodeling, lactate fermentation, and biosynthesis of octanoyl and myristate, and degradation of erythronate, threonate, galactitol, and allantoin were all significantly increased by KD3:1, decreased by KD4:1 and MCT2.5:1 (Supplementary Fig. 2d), and associated with low dietary fiber content (Supplementary Fig. 2e). The only pathway decreased by KD3:1, but elevated by KD4:1 and MCT2.5:1, was L-glutamate and L-glutamine biosynthesis (Supplementary Fig. 2d), which was further positively associated with dietary fiber (Supplementary Fig. 2e). Taken together, these results indicate that resistance vs. susceptibility to 6-Hz seizures in response to different KD infant formulas is associated with differential alterations in the composition and functional potential of the gut microbiome.

### Fiber content in the KD drives microbial alterations and promotes seizure resistance

The gut microbiome is shaped by changes in host diet and can be responsive to the presence, abundance, and sources of dietary macronutrients^39^. To gain insight into how different clinical KD formulas differentially alter the gut microbiome, we screened various dietary parameters for their effects on a model human infant microbial community. 9 bacterial strains were selected based on their prevalence and relative abundances across multiple large studies of the infant gut microbiome^40,41^ (**Supplementary** Fig. 3a**, Supplementary Data 3**). All community members were confirmed to grow stably together in a rich complex medium^42^ as a positive control (**Supplementary** Fig. 3b). To test the effects of KD fat ratio, the model infant gut microbial community was cultured in synthetic KD media prepared in ratios from KD4:1 to KD1.5:1 (**Supplementary** Fig. 3c**, Supplementary Data 6).** There were no statistically significant differences in taxonomic response to the KDs with different fat ratio (PERMANOVA, p=0.13, R^2^=0.14, **Supplementary** Fig. 3d). To examine effects of KD fat source, the model infant gut microbial community was cultured in synthetic media representing KD4:1, KD3:1, or MCT2.5:1, each using sunflower oil (6% saturated fat), soy lecithin (23% saturated fat and dominant in KD4:1 infant formula), or palm oil (50% saturated fat), as fat sources with different levels of saturation (**Supplementary** Fig. 3e). The media prepared with soy lecithin increased the absolute abundance of *B. infantis*, *B. fragilis*, and *C. perfringens*, resulting in distinct separation along PCoA1 from the sunflower and palm oil groups (PERMANOVA, p<0.05; **Supplementary** Fig. 3f). This may be due to the presence of free sugars (8%) in the commercial soy lecithin and/or the emulsifying properties of soy lecithin, compared to the other fat sources^43^. There were no statistically significant differences between the sunflower and palm oil groups across all media conditions **(Supplementary** Fig. 3f), suggesting that the differential effects of soy lecithin are driven by fat source rather than saturation level.

To test effects of additional fat sources, KD-based media were also prepared with addition of MCT, dominant in MCT2:5:1 infant formula, or linoleic acid, dominant in KD3:1 infant formula (**Supplementary** Fig. 3g). Addition of MCT increased the absolute abundance of *B. breve*, *B. infantis*, and *B. longum* compared to corresponding controls, resulting in notable shifts in diversity when added to KD4:1 and KD3:1 media (PERMANOVA p=0.05, R^2^=0.33; p=0.017, R^2^=0.32), but not KD2.5:1 media (PERMANOVA p=0.55, R^2^=0.04) (**Supplementary** Fig. 3h). In contrast, addition of linoleic acid decreased the absolute abundance of *B. infantis* and *B. vulgatus*, which resulted in statistically significant shifts across PCoA1 relative to all media groups (**Supplementary** Fig. 3h). This raises the question of whether differential effects of linoleic acid on the microbiome could contribute to the failure of KD3:1 infant formula to protect against 6-Hz seizures (**Fig. 1c, Supplementary** Fig. 1c).

Finally, to evaluate effects of carbohydrate type, the model infant gut microbial community was cultured in synthetic media representing KD4:1, KD3:1, and MCT2.5:1 and containing either lactose or a fiber mix, comprised of equal amounts of FOS, inulin, cellulose, and gum arabic, as the fiber sources that distinguish KD4:1 and MCT2.5:1 infant formula from KD3:1 and CD formulas (**Fig. 3a**). The presence of dietary fiber led to substantial shifts in the model infant gut microbial community across all media conditions, with particular enrichment of *B. fragilis* and decreases in *B. breve* and *B. infantis* (**Supplementary** Fig. 3i). PCoA analysis of synthetic metagenomic data assembled from quantitative taxonomic profiles showed notable clustering of fiber mix groups away from lactose controls (PERMANOVA, p=0.02 (KD4:1), p=0.08 (KD3:1), p=0.001 (MCT2.5:1), **Fig. 3b**), with greater discrimination than seen with alterations in fat ratio or source (**Supplementary** Fig. 3d-h). In particular, fiber mix yielded statistically significant decreases in several pathways related to amino acid biosynthesis, nucleotide and nucleoside biosynthesis, and carbohydrate degradation, among many others (**Fig. 3c and Supplementary** Fig. 3g). Among the 110 metagenomic pathways that were significantly altered by *in vitro* culture of the simplified infant microbial community with fiber mix compared to lactose **(Supplementary** Fig. 4), 15 pathways (13.6%) were similarly significantly altered in the fecal microbiome of mice fed the fiber-containing KD4:1 and MCT2.5:1, as compared to lactose-containing CD controls (**Fig. 3c**). Specifically, queuosine biosynthesis and its intermediate preQ_0_ biosynthesis were significantly enriched by fiber in the *in vitro* system and by fiber-containing KDs in the mouse. Similarly, fiber-induced decreases in pentose phosphate pathways, pathways related carbohydrate degradation (sucrose, glucose, xylose, and glycogen degradation), carbohydrate biosynthesis (UDP-N-acetyl-D-glucosamine biosynthesis and UDP-glucose derived O-antigen building blocks biosynthesis), amino acid biosynthesis (L-alanine, L-lysine and L-aspartate and L-asparagine biosynthesis), partial TCA cycle, and methylerythritol phosphate pathway were also shared with mouse metagenomes of KD4:1 and MCT2.5:1 groups (**Fig. 3c and 2e**). The results suggest that dietary fiber, more so than fat ratio or source, exerts a strong influence on community structure and functional potential of a model infant gut microbial community. Select alterations are consistent with those seen in the mouse microbiome in response to host consumption of fiber-containing clinical KD infant formulas (KD4:1 and MCT2.5:1), which confer resistance to 6-Hz seizures (**Supplementary** Fig. 4). The results suggest that these particular metagenomic signatures may serve as biomarkers for seizure resistance.

**Figure 3.**
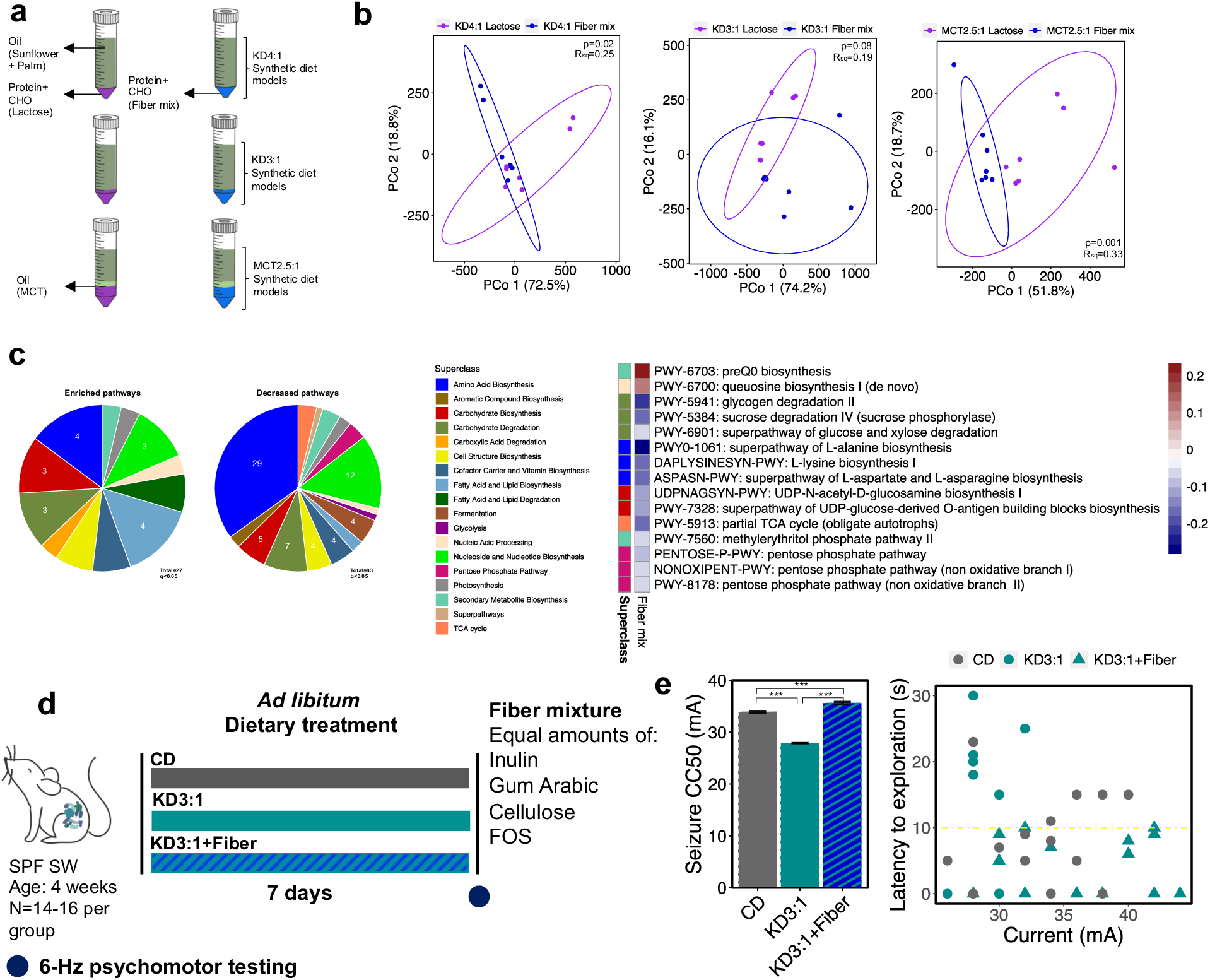
Addition of dietary fiber to KDs enriches metagenomic features associated with seizure protection in a model human infant gut community and restores resistance to 6-Hz seizures in mice. a. Experimental design: Fiber mix containing inulin, gum arabic, cellulose, and fructooligosaccharide (FOS), or lactose as a non-fiber carbohydrate control, was added to KD-based synthetic culture media for anaerobic culture of a model human infant gut microbial community b. Principal coordinates plots of metagenomic pathway abundance data for human infant microbes grown in KD-based media containing fiber mix versus lactose. (PERMANOVA, n=7/condition) c. Venn diagram of differential metagenomic pathways (q<0.05) shared across all fiber-containing KD media groups relative to corresponding lactose-containing media groups as controls (left). 15 fiber-induced differential metagenomic pathways (q<0.05) that are similarly seen in seizure protective mice fed KD4:1 or MCT2.5:1 (right). (General Linear Model, n=7/condition) d. Experimental design: 4 week old conventional (specific pathogen free, SPF) Swiss Webster (SW) mice (n=14-16 mice/group) were fed KD3:1 supplemented with fiber mix, KD3:1 alone, or CD as liquid diets for 7 days. e. 6-Hz seizure threshold (left) and latency to exploration (right) for mice fed KD3:1+fiber mix, KD3:1, or CD as liquid diet (left, one-way ANOVA with Bonferroni, n=14-16 mice/group, ***p<0.001). Yellow line at y = 10 s represents threshold for scoring seizures.

To test whether dietary fiber content has a causal impact on resistance to 6-Hz seizures, we supplemented the fiber mix into the KD3:1 infant formula to match reported fiber levels in KD4:1 infant formula, and tested mice for seizure susceptibility at 7 days after dietary treatment (**Fig. 3d**). As previously demonstrated, mice fed liquid KD3.1 exhibited decreased seizure threshold compared to CD controls (**Fig. 3e**). Notably, addition of fiber to the KD3:1 elevated seizure thresholds to levels that exceeded those seen in CD controls. We further repeated the fiber supplementation using the solid diet paradigm, where the same infant formulas were dehydrated and administered as chow instead of liquid diet. As seen in liquid form, supplementation with fiber mix significantly increased seizure threshold of mice fed KD3:1, with no significant differences in diet consumption (**Supplementary** Fig. 5). These data demonstrate that addition of fiber to the low fiber KD3:1 infant formula restores its antiseizure effects toward levels seen with fiber-containing KD4:1 and MCT2.5:1.

To determine whether dietary fiber supplementation can potentiate KD-induced seizure protection, we supplemented the fiber-containing KD4:1 infant formula, which yielded the highest seizure thresholds of all KD variants (**Fig. 1**), with the dietary fiber mix that is already existing in the formula and tested mice for resistance to 6-Hz seizures after 7 days of feeding with the liquid diet (**Fig. 4a**). The additional fiber added to KD4:1 formula increased fiber content from 5.3% to ∼10.3%. Dietary fiber supplementation significantly increased seizure thresholds to levels that exceeded those seen with KD4:1 alone (**Fig. 4b**). There were no significant differences between groups in dietary consumption (**Supplementary** Fig. 6a). The ability of fiber supplementation to further promote the anti-seizure effects of KD4:1 was similarly seen when administered as solid diet, instead of liquid diet, also with no significant differences in food consumption (**Supplementary** Fig. 6b, c). Short-chain fatty acids (SCFAs) are primary end products of gut microbial fiber fermentation in the colon and have been shown to impact host brain activity and behavior^44^. To further ask whether fiber supplementation promotes seizure resistance via SCFAs, we supplemented KD4:1 infant formula with the SCFAs acetate, butyrate, and propionate, at concentrations predicted to match those achieved produced by fermentation of the dietary fiber mix. In both liquid and solid form, SCFA supplementation failed to phenocopy effects of dietary fiber supplementation and instead yielded mice with modest reductions in resistance to 6-Hz seizures, as compared to controls supplemented with vehicle solution (**Supplementary** Fig. 7a, b). Taken together, these data indicate that dietary fiber supplementation both restores the anti-seizure effects of the low fiber KD3:1 and further potentiates the anti-seizure effects of the fiber-containing KD4:1, through mechanisms that are not recapitulated by oral SCFA supplementation.

**Figure 4.**
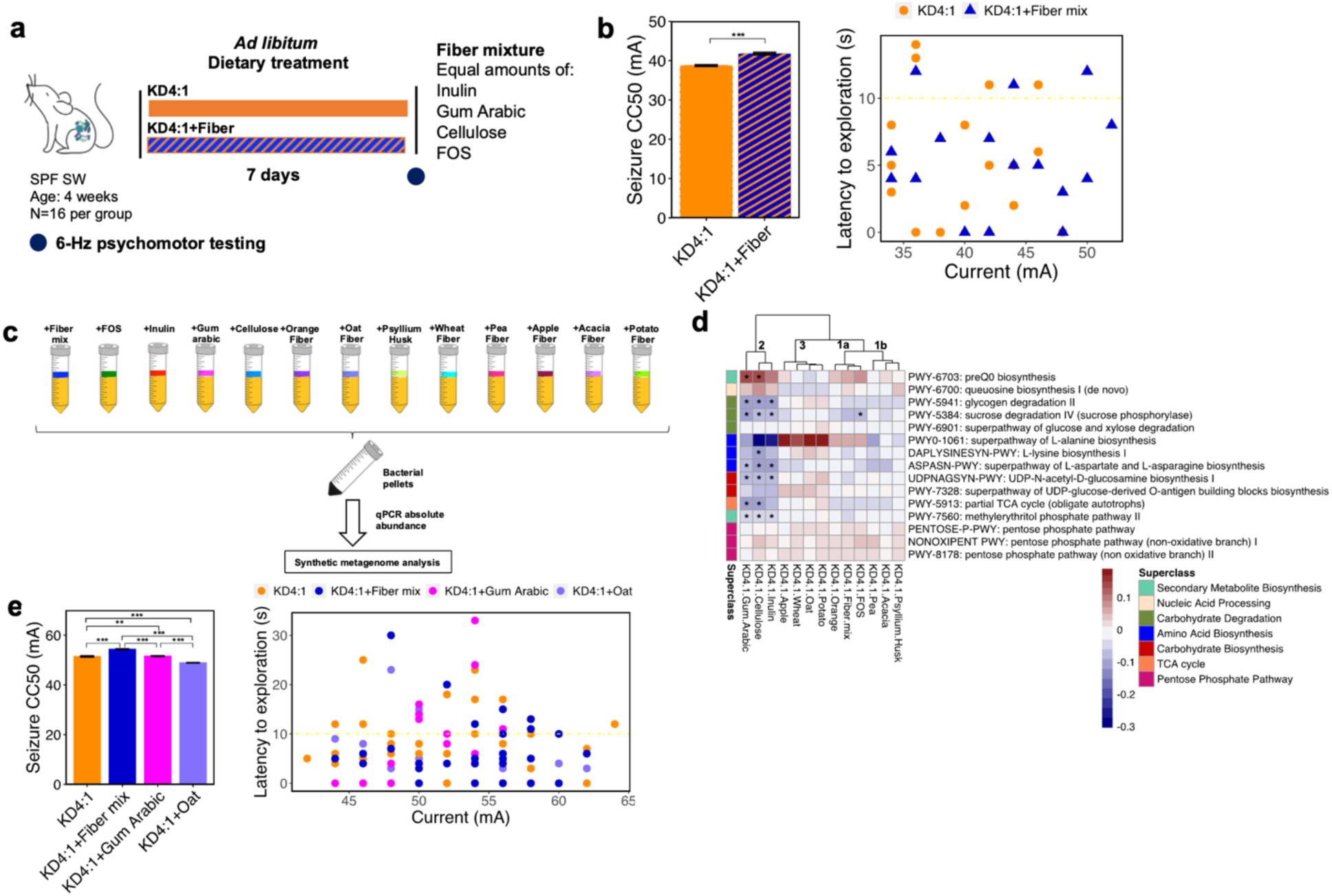
Addition of excess dietary fiber to fiber-containing KD4:1 further potentiates seizure resistance. a. Experimental design: 4 week old conventional (specific pathogen free, SPF) Swiss Webster (SW) mice (n=16 mice/group) were fed KD4:1 supplemented with fiber mix or KD4:1 alone as liquid diets for 7 days. b. 6-Hz seizure threshold (left) and latency to exploration (right) for mice fed KD4:1 and KD4:1+fiber mix as liquid diet (left, Welch’s t-test n=16 mice/group, ***p<0.001). Yellow line at y = 10 s represents threshold for scoring seizures. c. Experimental design: 13 dietary fiber sources and types were supplemented to KD4:1 infant formula for anaerobic culture of a model human infant gut microbial community d. Heatmap of 15 fiber-induced differential metagenomic pathways (q<0.05) that were similarly seen in seizure-protected mice fed KD4:1 or MCT2.5:1 (right). Groupings were denoted on top of the dendrogram. (General Linear Model statistical test, n=8-10/condition, * q<0.05 for fiber source/type relative to KD4:1 as a control) e. 6-Hz seizure threshold (left) and latency to exploration (right) for mice fed KD4:1 supplemented with dietary fiber mix (Group 1), gum arabic (Group 2), or oat fiber (Group 3), or KD4:1 alone as paste diet (left, one-way ANOVA with Bonferroni, n=14 mice/group, **p<0.01, ***p<0.001). Yellow line at y = 10 s represents threshold for scoring seizures.

### Different fiber types and sources elicit differential microbial alterations and seizure outcomes

Dietary fibers are fermented by select gut bacteria and shape the composition and activity of the gut microbiome^45^. To gain insight into whether particular fiber types or sources interact with KD4:1 to differentially alter the infant gut microbiome, we screened 13 different fiber conditions, comprised of commercially available fiber products or purified fiber types, for their additional effects on the model infant microbial community when grown directly in KD4:1 infant formula (rather than in a diet-based synthetic culture medium, as in prior experiments) (**Fig. 4c**). Taxonomic profiles showed that 8 out of the 13 fiber conditions significantly increased the absolute abundance of *B. fragilis*, and 11 fiber conditions significantly decreased *B. breve* (**Supplementary** Fig. 8), both of which align with previous *in vitro* results from fiber supplementation into synthetic media (**Supplementary** Fig. 3i). 7 of the 13 fiber conditions yielded reductions in *E. coli*, which parallel the increases in *E. coli* observed with mouse consumption of fiber-deficient KD3:1 (**Supplementary** Fig. 2c). We next generated synthetic metagenomic profiles for the 13 fiber supplementation conditions and filtered results to prioritize the 15 protective features that were shared between mouse consumption of the KD4:1 and MCT2.5:1 (**Fig. 2e**) and model human infant microbial community responses to fiber in synthetic diet-based media (**Fig. 3c, Supplementary** Fig. 4). The results revealed 4 subgroupings of model infant microbial responses to the 13 different fibers in KD4:1 infant formula (**Fig. 4d)**. Group 1a consisted of fiber mix, FOS, and orange fiber and was characterized by increases in genes related to preQ0 biosynthesis and L-alanine biosynthesis, with reductions in sucrose degradation and partial TCA cycle (**Fig. 4d**). Group 1b consisting of pea, acacia, and psyllium husk fibers, clustered together with Group 1a and exhibited a similar general pattern of metagenomic features but with reductions in L-alanine biosynthesis and less substantial shifts in preQ0 biosynthesis and sucrose degradation (**Fig. 4d**). Group 2 consisted of inulin, cellulose, and gum arabic, which was characterized by significant decreases in genes related to 5-7 pathways (glycogen and sucrose degradation, L-alanine, L-lysine, L-aspartate, L-asparagine, and UDF-N-acetyl-D-glucosamine biosynthesis, partial TCA cycle, and methylerythritol phosphate pathway) and significant increases in preQ0 biosynthesis genes (**Fig. 4d**). Group 3, consisting of oat, potato, wheat, and apple fibers, was characterized by notable increases in representation of L-alanine biosynthesis and UDP-glucose-derived O-antigen building blocks biosynthesis, with decreases in queuosine biosynthesis (**Fig. 4d**).

Based on these patterns of microbial representation for key metagenomic features conserved in mice fed fiber-containing KDs and infant microbial communities cultured with fiber-supplemented media, we selected one representative fiber condition per primary grouping (Group 1: fiber mix, Group 2: gum arabic, Group 3: oat fiber) to test for causal effects on seizure resistance. We supplemented representative fibers from each group into KD4:1 infant formula to raise fiber content from 5.3% to ∼10.3%, and tested mice for resistance to 6-Hz seizures at the 7^th^ day after feeding in paste form. As previously observed in liquid and solid diet form (**Fig. 4b, Supplementary** Fig. 5b), supplementation of KD4:1 paste with fiber mix significantly increased resistance to 6-Hz seizures (**Fig. 4e**). In contrast, supplementation with gum arabic (Group 2) had no overt effects on seizure threshold compared KD4:1 controls (**Fig. 4e**). In addition, supplementation with oat fiber (Group 3) had a detrimental effect, significantly decreasing seizure thresholds compared to KD4:1 controls and all other fiber conditions (**Fig. 4e**). Overall, these data reveal that the ability of fiber supplementation to potentiate the seizure protective effects of KD4:1 infant formula is specific to particular sources and types of fibers that alter key metagenomic features of the gut microbiome.

## Discussion

Findings from this study demonstrate that different clinical KD infant formulas have varying effects on seizure resistance in mice, likely due to differences in how specific dietary components affect the function of the gut microbiome. We find that fiber-containing commercial infant formulas KD4:1 and MCT2.5:1 promote resistance to 6-Hz seizures in mice, whereas the fiber-deficient commercial infant formula KD3:1 increases susceptibility to 6-Hz seizures. Correspondingly, the protective KD4:1 and MCT2.5:1 induce several shared metagenomic alterations in the gut microbiome, which are not seen with KD3:1. In particular, KD4:1 and MCT2.5:1, but not KD3:1, reduce representation of select genes related to carbohydrate degradation, which were significantly associated with the presence of dietary fiber and similarly induced by fiber supplementation to a cultured infant gut microbial community. Adding a fiber mixture to the KD3:1 to match levels present in KD4:1 and MCT2.5:1 restores seizure protection in mice. Moreover, supplementing the fiber mixture to the already protective KD4:1 infant formula further enhances seizure resistance in mice.

Only a few small human studies have tested the effects of different medical KD regimens on seizure reduction, reporting no significant differences between the MAD, MCT, and LGIT diets relative to the classic KD in controlling seizures in children with refractory epilepsy^25,26,28,29^. However, none of these examined the role of fiber or any specific dietary constituents on patient responses to KD therapy. A cross-sectional study of 150 epileptic individuals reported insufficient intake of fiber, among several other vitamins and minerals, and that patients with low intake of vegetables exhibited greater likelihood of uncontrolled seizures^46^. When considering specific macronutrients and micronutrients that distinguish patients with controlled and uncontrolled seizures, percent intake of fiber was the closest to statistical significance (reported p=0.05). In addition, a human study of KD therapy in children with refractory epilepsy reported changes in 29 metagenomic pathways, including the reduction of seven pathways involved in carbohydrate metabolism and fermentation such as fructooligosaccharides (FOS) and raffinose utilization, sucrose utilization, glycogen metabolism, lacto-N-biose I and galacto-N-biose metabolic pathway; lactate, pentose phosphate pathway; and formaldehyde assimilation: ribulose monophosphate pathway^16^. Further research is needed to explore the role of dietary fiber and specific dietary nutrients on the clinical efficacy of KD formulations for treating refractory epilepsy.

Dietary fibers are resistant to digestion by the host and specifically fermented by gut bacteria that together encode hundreds of glycoside hydrolases with varying specificity for different fiber types^47^. As such, not only does the gut microbiome degrade fiber, it also responds to and is shaped in composition and function by dietary fiber. We find that supplementing mice with the SCFAs butyrate, propionate, and acetate, as common microbial end-products of fiber fermentation, fails to phenocopy the beneficial effects of fiber supplementation on potentiating seizure protection in mice fed the KD4:1. This may align with prior human studies reporting that epilepsy is associated with deficient levels of SCFA-producing bacteria, which are further reduced by KD therapy to promote seizure control^16,22,48^. Beyond SCFAs, several other carboxylic acid metabolites, neurotransmitters, vitamins, and bile acids are also modulated by fiber fermentation^49,50^. This suggests that the fiber effects on seizure resistance may not be mediated by common SCFAs , but rather by other non-SCFA metabolites generated by fiber fermentation or indirect effects of fiber fermentation on the microbiome and host. Indeed, alterations in the gut microbiome are increasingly implicated in risk for epilepsy and seizure responsiveness to the KD across several humans studies^14–19^.

Findings from animal models establish proof-of-principle that KD-induced alterations in the gut microbiome contribute to seizure resistance^12,20–23^, suggesting that differential effects of dietary formulations on the gut microbiome may lead to variation in seizure protection. By screening various dietary factors that distinguish the KD infant formulas, including fat ratio, fat source, fat saturation, and carbohydrate type, on a model human infant microbial community, we find that addition of fiber to a diet-based synthetic culture media elicits substantial shifts in microbial metagenomic profiles. Many key metagenomic features seen in response to fiber supplementation in the *in vitro* system are consistent with those seen in the gut microbiome of seizure-protected mice fed fiber-containing KD4:1 and MCT2.5:1, suggesting direct interactions between dietary fiber and the microbiome that are effectively modeled in simplified microbial culture systems.

Different fiber types and sources can vary greatly in their chemical structure, fermentability, and effects on the gut microbiome^51^. We further expanded our *in vitro* screening approach to include 13 different soluble or insoluble fiber types and sources, as supplemented directly into the commercial KD4:1 infant formula (rather than a diet-based synthetic culture medium). By using key fiber-associated metagenomic features to stratify microbial responses to the 13 fiber conditions, we identified a specific subset of fibers that potentiate the seizure protective effects of the KD4:1 in mice. This subgroup, including fiber mix (inulin, FOS, gum arabic, cellulose), FOS alone, and orange fiber, is characterized by metagenomic alterations in pathways related to preQ0 biosynthesis, L-alanine biosynthesis, sucrose degradation, partial TCA cycle. PreQ_0_ is a deazapurine nucleoside with reported antibiotic, anticancer, antineoplastic, and antiviral properties^52,53^. In mice that exhibited seizure resistance in response to transplantation of the clinical KD-induced human microbiota, microbial preQ_0_ biosynthesis was associated with alterations in hippocampal expression of genes related to neuron generation and migration protection^20^. L-alanine is an essential amino acid that is modulated by ketosis^54^ and regulates the function of glutamatergic neurons and astrocytes^55^. L-alanine levels were diminished significantly in the cerebrospinal fluid of children after four months of KD therapy^56^, and genes related to L-alanine metabolism were elevated in imputed microbial metagenomic pathways from epileptic individuals relative to healthy controls^57^. Alterations in L-alanine biosynthesis could result in differential levels of 2-oxoglutarate, which is involved in the production of pyruvate and glutamate. Glutamate biosynthesis was positively associated with the presence of dietary fibers in clinical KDs and negatively associated with fiber-deficient seizure susceptible group KD3:1 **(Supplementary** Fig. 2d,e). In bacteria, the TCA pathway fuels aerobic respiration, wherein acetyl-CoA is converted to intermediate organic acids such as citrate, 2-oxoglutarate, and succinate. Specifically, 2-oxoglutarate also known as α-ketoglutarate is an important intermediate in TCA cycle and is known to affect glutamate and GABA levels in the brain^58^. Similarly, the dietary intake of 2-oxoglutarate (α-ketoglutarate) decreases the α-synuclein pathology in mouse model of Parkinson’s disease^59^. Even though this metabolite has been implicated in neurophysiological conditions, how microbial levels of 2-oxoglutarate and other intermediate metabolites from the TCA cycle, may affect the brain to alter anti-seizure susceptibility is unknown. During KD therapy, patients supplemented with oral citrate as an alkalizing agent prevented metabolic acidosis without affecting the 7-month efficacy rates^59^. Similarly, KD has been shown to affect the succinate levels through succinate dehydrogenase activity in rodent models of aging^60^ and through effects on mitochondrial respiration to restore ATP production^61^. Microbial sucrose utilization is a carbohydrate pathway reduced after KD therapy in humans^16^, likely due to the low availability of carbohydrates in the diet. This specific pathway, sucrose degradation IV, is mainly encoded by *Bifidobacterium* species and shunts β-D-fructofuranose-6-phosphate to produce acetate, lactate, formate, and acetyl-coA^62^, which aligns observed fiber-induced reductions in *B. breve* **(Supplementary** Fig. 8). Overall, these results suggest that increases in microbial biosynthetic pathways for preQ_0_ and L-alanine and reductions in microbial carbohydrate metabolism may serve as biomarkers for diet-induced seizure resistance. Further research is needed to determine whether there are causal links between these particular microbial functions and seizure protection.

Altogether, results from this study reveal that nuanced differences in the formulation of KDs that are used to treat refractory epilepsy can lead to major differences in treatment efficacy and in the functional potential of the gut microbiome. In particular, we highlight a major role for dietary fiber in restoring and potentiating the seizure protective effects of commercial KD formulas when fed to mice. We demonstrate that dietary fiber shifts key metagenomic features in both the mouse gut microbiome and a model human infant microbial community, which can be used to identify specific fiber types that potentiate the seizure protective effects of a classical KD formula in mice. Our findings align with increasing evidence that the gut microbiome modifies the anti-seizure effects of the KD and that microbiome-targeted diets can be used to shape the structure and function of the gut microbiome. It further supports the growing notion that careful consideration of dietary effects on host-microbial interactions is needed to inform the design of more effective and personalized dietary interventions for disease.

## Methods

### Mice

All mouse experiment protocols were approved by the UCLA Institutional Animal Care and Use Committee. Juvenile (4-week old) specific pathogen free (SPF), male Swiss Webster (Taconic Farms) mice were used for all animal experiments, fed standard chow (Labdiet 5010, 28.7%: 13.1%: 58.2% protein: fat: carbohydrate by calories), and housed in sterile caging under a 12 h:12 h light:dark cycle with standard temperature and humidity control.

### Dietary treatment

Experimental animals were fed commercially available KD infant formulas (KetoCal, Nutricia North America, Fig. 1a, Supplementary Data 1) or a popular commercially available, standard infant formula as control diet (Abbott Nutrition, Fig. 1a, Supplementary Data 1) for 7 days. For liquid diet paradigm, 90 g of powder formula or 90 mL of liquid formula was mixed with 600 mL water at 60°C. Before adding to cages, the diet solution was brought to 1L and each cage containing 3-4 mice was supplemented with liquid diets in water bottles. The water bottles were filled with liquid diets and the cages were changed every 1-2 days. For the solid diet paradigm, 90 g of powder formula was mixed with 600 mL water and dehydrated using a food dehydrator (CASORI). The diets were administered in sterile petri dishes and cages were provided with standard sterile water. For the pasted diet paradigm, 30 g of powder was mixed with water and administered as a paste in sterile petri dishes.

For fiber supplementation experiments, 5 g of individual fiber or fiber mixture (fructooligosaccharides (FOS), inulin, cellulose and gum arabic from Sigma-Aldrich, mixed at 1:1 (w/w)) was added to 90 g of KD formula prior to administering as a paste as described above. For SCFA supplementation, we considered the following concentrations as reference values for SCFAs reported in SPF mice fed standard chow containing 15% fiber: acetate (67.5 mM), propionate (25 mM) and butyrate (40 mM)^63^. To model 5% fiber content present in KD4:1, we therefore administered sodium acetate (22.55 mM), sodium propionate (8.33 mM) and sodium butyrate (13.35 mM) in sterile drinking water. For paste diets, 1:10 of the SCFA mixture were mixed with water and added to the powder diets at the following concentrations: sodium acetate (2.255 mM), sodium propionate (0.833 mM) and sodium butyrate (1.335 mM). As a negative control, sodium chloride (NaCl) was supplemented to match amounts in SCFA salts in water (132.5 mM) and in diet (13.25mM).

### 6-Hz psychomotor seizure assay

The 6-Hz psychomotor seizure test was conducted as previously described^12^. One drop (∼50 ul) of 0.5% tetracaine hydrochloride ophthalmic solution was applied to the corneas of each mouse 10-15 min before stimulation. Corneal electrodes were coated with a thin layer of electrode gel (Parker Signagel). A current device (ECT Unit 57800, Ugo Basile) was used to deliver current at 3 s duration, 0.2 ms pulse-width and 6 pulses/s frequency. CC50 (the intensity of current required to elicit seizures in 50% of the experimental group) was measured as a metric for seizure susceptibility. Pilot experiments were conducted to identify 28 mA as the CC50 for SPF wild-type Swiss Webster mice when they are on liquid and solid diet and 44 mA when they are on paste diet. Each mouse was seizure-tested only once, and thus n=14-16 mice were used to adequately power each experimental group. 28 or 44 mA currents were administered to the first mouse per cohort, followed by fixed increases or decreases by 2 mA intervals. Mice were restrained manually during stimulation and then released into a new cage for behavioral observation. Locomotor behavior was recorded using a camera and quantitative measures for stunned fixture, falling, tail dorsiflexion (Straub tail), forelimb clonus, eye/vibrissae twitching, and behavioral remission were scored manually. Latency to exploration (time elapsed from when an experimental mouse is released into the observation cage (after corneal stimulation) to its normal exploratory behavior) was scored manually with an electronic timer. Mice were blindly scored as protected from seizures if they did not show seizure behavior and resumed normal exploratory behavior within 10 s. Seizure threshold (CC50) was determined as previously described^64^, using the average log interval of current steps per experimental group, where sample n is defined as the subset of animals displaying the less frequent seizure behavior. Data used to calculate CC50 are also displayed as latency to explore for each current intensity, where n represents the total number of biological replicates per group regardless of seizure outcome.

### Fecal shotgun metagenomics

Frozen stool samples from mice pre- and post-dietary treatment were subjected to DNA extraction using the ZymoBIOMICS DNA Miniprep kit (Zymo), with bead beating used to lyse cells. Briefly, the samples were transferred into PowerBead tubes containing lysis solution and bead beaded at maximum speed for 1 min five times with 1 min of ice incubation in between cycles. The rest of the protocol followed the manufacturer’s instructions. The DNA was eluted in 60 μL elution buffer provided by the kit. Purified DNAs were sent to Novogene Corporation Inc for paired end (PE) metagenomic sequencing. Sequencing was performed on the Illumina NovaSeq platform with PE reads of 150 bp for each sample averaging around 3GB data. Raw reads were subjected to kneaddata to remove host contaminants. Metagenomic data was analyzed using HUMAnN3^65^ and MetaCyc database to profile gene families and pathway abundance. MetaPhlAn4 was used for metagenomic taxonomic profiling^66^. α-diversity indexes for taxonomic profiling were determined by Shannon’s index, richness, and Pielou’s Evenness using vegan v2.6-4 in R. For β- diversities, calculate_diversity.R script were run within the MetaPhlAn4. For Unifrac distances, mpa_vOct22_CHOCOPhlAnSGB_202212.nwk was used for SGB-level phylogenetic tree as reference. R packages tidyverse v2.0.0, vegan v2.6-4, and phyloseq v1.38.0 was used for Principal coordinate analysis (PCoA) of taxanomic distribution. Alterations in microbial diversity were assessed using PERMANOVA with adonis2 with 999 permutations from the vegan package in R. File2meco R package was used for MetaCyc pathway hierarchical classification^67^. MaAsLin 2.0^68^ was used to assess significant pathway associations between dietary treatments with an adjusted p value (q value) cutoff of 0.05, where indicated in the figure by asterisk.

### Beta-hydroxybutyrate (BHB) measurements

Blood was collected via a capillary tube from the medial canthus of the eye, allowed to clot 30 min at room temperature, and spun through SST vacutainers (Becton Dickinson) at 1500g for 90 sec for serum separation. Samples were immediately snap frozen in liquid nitrogen and stored at - 80°C until further processing. BHB levels were quantified by colorimetric assay according to the manufacturer’s instructions (Cayman Chemical).

### Bacterial strains and culturing

The following strains were selected to represent the human infant gut microbiome, based on their high relative abundance in their respective phyla and prevalence at >1% relative abundance across the study population^40,41^ (**Supplementary** Fig. 3a). Type strains were obtained either from ATCC or DSMZ collection and propagated as instructed: *Bifidobacterium longum* subsp. *infantis* DSM 20088, *Bifidobacterium longum* subsp. *longum* ATCC BAA-999, *Bifidobacterium breve* DSM 20213, *Bacteroides fragilis* ATCC 25285, *Bacteroides vulgatus* ATCC 8482, *Enterococcus faecalis* ATCC 19433, *Clostridium perfringens* ATCC 13124, *Escherichia coli* K-12 ATCC 10798, *Klebsiella pneumoniae* subsp. *pneumoniae* ATCC 13883. The cultures were routinely grown anaerobically in their respective media and temperature (**Supplementary Data 3**). The growth of species were tested on a rich complex medium^42^ for 24 h to confirm stable relative abundances over the duration of anaerobic culture, as confirmed by cfu plating and qPCR (**Supplementary Data 2, Supplementary** Fig. 3b).

### *In vitro* batch culture fermentations

Synthetic KDs with different ratios, fat and carbohydrate source were prepared using sunflower oil (Baja Precious), vegetable shortening (Crisco), palm oil (Ökonatur), soy lecithin (Modernist Pantry), linoleic acid (Sigma-Aldrich), and Medium Chain Triglycerides (MCT, Nutriticia) as fat sources, whey protein isolate (Bulk Supplements) as protein source, and lactose (modernist pantry) and dietary fiber mixture of fructooligosaccharides (Sigma-Aldrich), inulin from chicory (Sigma-Aldrich), crystalline cellulose (Sigma-Aldrich), and gum arabic from Acacia Tree (Sigma-Aldrich) as carbohydrate sources. Additionally, for fiber supplementation fermentation experiments, wheat, pea, potato, and apple fiber from J. Rettenmaier USA LP, orange (citrus) fiber from Citri-Fi Naturals, oat (NuNaturals), acacia (Nutricost organic), and psyllium husk (It’s just) were used. The powders were ultraviolet (UV)-sterilized and confirmed to be sterile by aerobic and anaerobic culture. They were then mixed with simulated saliva solution, gastric solution, and intestinal fluid as described in INFOGEST model^69^ without enzymatic solution (**Supplemental Data 5**) to simulate the gastric and intestinal bolus entering to the colon and was subjected to an *in vitro* batch culture fermentation. The representative bacterial strains were mixed in a minimal media at the dilution factor (1:100) needed to achieve a ratio of 21% Actinobacteria, 14% of Bacteroidetes, 28% of Firmicutes, and 37% of Proteobacteria, reflective of relative abundances seen in a typical infant gut^40,41^ (**Supplemental Data 3**). Species that comprise Actinobacteria, Bacteroidetes and Firmicutes were mixed at 1:1 ratio, whereas Proteobacteria consists of 57 % of *Escherichia coli* and 42% of *Klebsiella pneumoniae*. The bacterial mixture was then mixed with each diet bolus (1:1 v/v) and subjected to 24-hour anaerobic culture. After 24 hours, the bacterial pellets were separated from the media and stored at -80°C until further analysis. The pellets from pre- fermentation were also collected as a control.

### Bacterial quantification via qRT-PCR

Total DNA was extracted from the pellets collected after fermentation, following standard procedures for the ZymoBIOMICS DNA Miniprep kit. The microbial composition was determined using quantitative RT-PCR with species specific primers^70–73^ and respective qPCR conditions (**Supplementary Data 5**). DNA extracted from individual overnight cultures were used to generate a standard curve. The copy numbers for each sample were calculated based on the standard curve and normalized to DNA concentration of the original sample. Absolute quantification of growth after anaerobic culture of each sample was determined by subtracting the pre-fermentation quantities and presented as log values. Any species that exhibited negative values after subtraction were regarded as zero or no growth. Data are presented in bar plots as a mean of each bacteria. PCoA plots were created using cmdscale from the distance matrix created using Euclidean distances in vegan package in R.

### Production of synthetic metagenome reads and synthetic metagenome analysis

The genome fastq files for each species were obtained from ATCC.org. Open source BBMap v38.94 randomreads.sh plugin was used to randomly produce paired reads at 150 bp length from each genome based on the qPCR absolute quantification multiplied by a million. For each sample, between 50-80 million metagenomic reads were produced. Metagenomes were analyzed using Humann3 and significant pathway associations were determined with MaAsLin2 package in R as described above.

### Statistical analysis

All statistical analyses were conducted using R version 4.1.2. Data for box-and-whisker plots were plotted as median with first and third quartiles. Data for parametric data sets was analyzed using one-way ANOVA with Bonferronii adjustment between multiple groups. For differences between two sample conditions, parametric datasets were analyzed using Welch’s t-test and non-parametric data sets using Wilcoxon signed rank test. For non-parametric distributions with more than two groups, data was analyzed by Kruskal-Wallis with Dunn’s test. For PCoA plots, the distance matrix created within vegan package was initially subjected to betadisper and permutest for multivariate homogeneity of groups dispersions (variances), then PERMANOVA with adonis2 with 999 permutations was used to determine statistical differences between groups. Significant differences from the tests were donated as follows: * p<0.05, **p<0.01, ***p<0.001, ****p<0.0001. Notable non-significant differences were denoted as n.s.

### Data Availability

Raw and processed data from metagenomic profiling from mice, metagenomic data created synthetically, and associated metadata are presented in Tables S7, S8, S9, and S10 and are available online through the NCBI Sequence Read Archive (SRA) repository at SRA: SUB14608643 XXXX

## Acknowledgments

We thank Joyce H. Matsumoto and Beck Reyes at the UCLA’s Mattel Children’s Hospital for valuable discussion on clinical KD treatments and the UCLA Human Nutrition Phytochemical Core Facility for technical support with LC-MS. This work was supported by NINDS grant #R01NS115537 (E.Y.H).

## Author Contributions

E.Ö, K.B.Y, G.R.L, J.P., and E.Y.H conceptualized and planned bacterial and mouse experiments. E.Ö, K.B.Y, J.H., L.D., and K.L. performed bacterial experiments. E.Ö, K.B.Y, and G.R.L performed mouse experiments. E.Ö performed metagenomics and analyzed data. E.Ö and E.Y.H wrote the manuscript. All authors contributed to final manuscript.

---

**Supplementary Figure 1.**
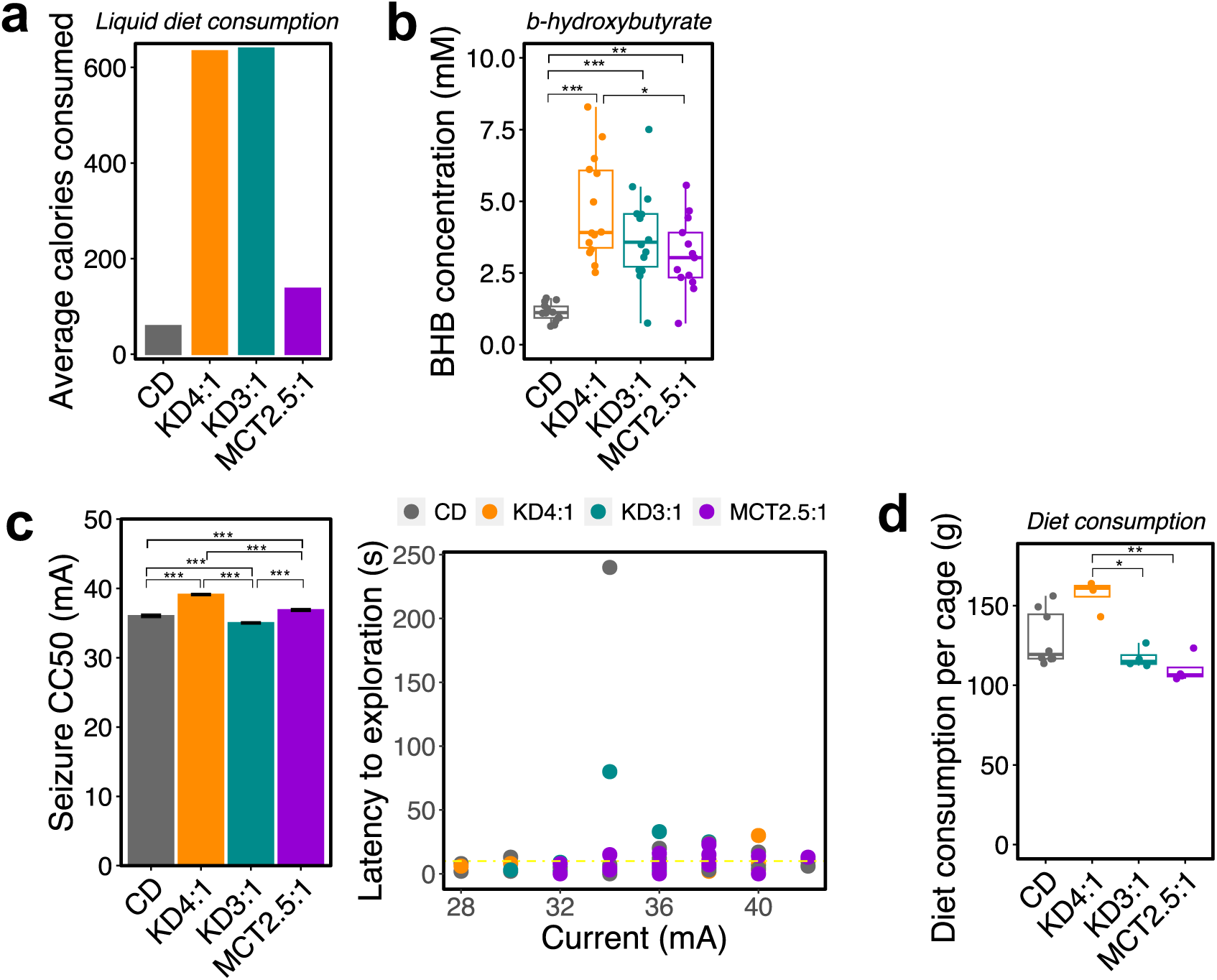
Medical KDs administered as solid diets phenocopy differential seizure responses seen with liquid diets. a. Average caloric intake per cage for KDs and CD administered as liquid diet (n=3-4 cages) b. Serum beta-hydroxybutyrate from mice fed liquid KDs or CD. (One way ANOVA with Bonferroni: ^∗^p < 0.05, **p<0.01, ***p<0.00; n=14 mice/group. Data are presented as box-and-whisker plots with median and first and third quartiles). c. 6-Hz seizure threshold (left) and latency to exploration (right) for mice fed KDs or CD as solid diet (left, one-way ANOVA with Bonferroni, n=16 mice/group, ***p<0.001). Yellow line at y = 10 s represents threshold for scoring seizures. d. Average consumption of solid diets (n=4 cages; Kruskal-Wallis with Dunn’s test: ^∗^p < 0.05, **p<0.01. Data are presented as box-and-whisker plots with median and first and third quartiles).

**Supplementary Figure 2.**
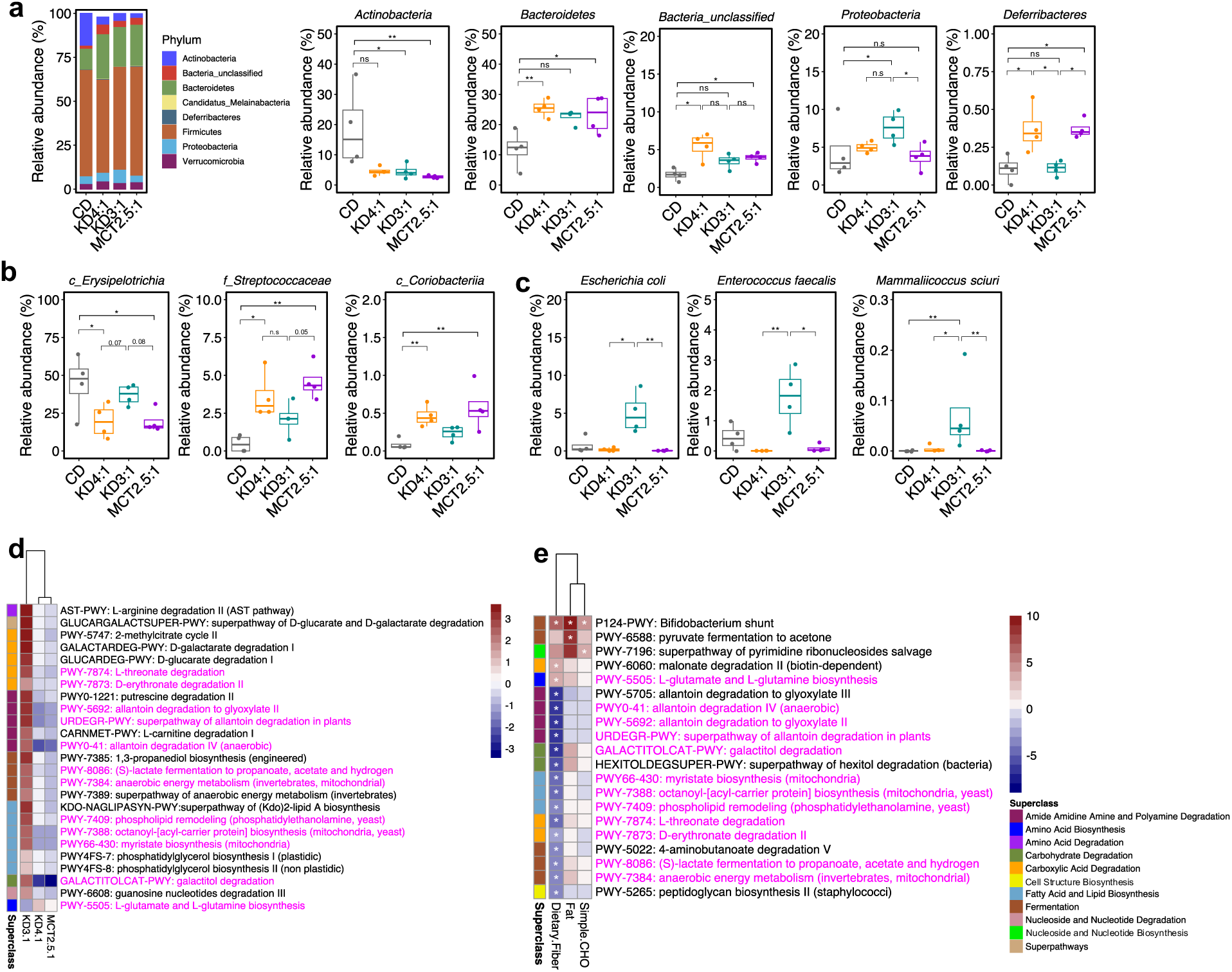
Effects of KDs on taxonomic and metagenomic signatures of the fecal microbiome in mice. a. Taxonomic distributions of bacterial phyla from fecal metagenomics data of mice fed liquid KDs or CD (left, n = 4 cages/group). Relative abundances of *Actinobacteria, Bacteroidetes, Bacteria_unclassified, Proteobacteria, and Deferribacteres* (right, n = 4 cages/group. Kruskal-Wallis with Dunn’s test: ^∗^p < 0.05, **p< 0.01 n.s., not statistically significant) b. Relative abundances bacterial taxa differentially altered by KD4:1 and MCT2.5:1, but not KD3:1 relative to CD. (n=4 cages/group. Kruskal-Wallis with Dunn’s test. ^∗^p < 0.05, **p< 0.01, n.s., not statistically significant. Data are presented as box-and-whisker plots with median and first and third quartiles). c. Relative abundances of bacterial taxa differentially altered by KD3:1, but not KD4:1 and MCT2.5:1, relative to CD. (n=4 cages/group. Kruskal-Wallis with Dunn’s test. ^∗^p < 0.05, **p< 0.01. Data are presented as box-and-whisker plots with median and first and third quartiles). d. Heatmap of differential metagenomic pathways (q<0.05) seen in seizure susceptible group KD3:1, but not seizure protective groups KD4:1 and MCT2.5:1. (General Linear Model, *q<0.05, n=4/condition) e. Heatmap of metagenomic pathways that are significantly associated with macronutrient composition (General Linear Model, *q<0.05, n=4/condition)

**Supplementary Figure 3.**
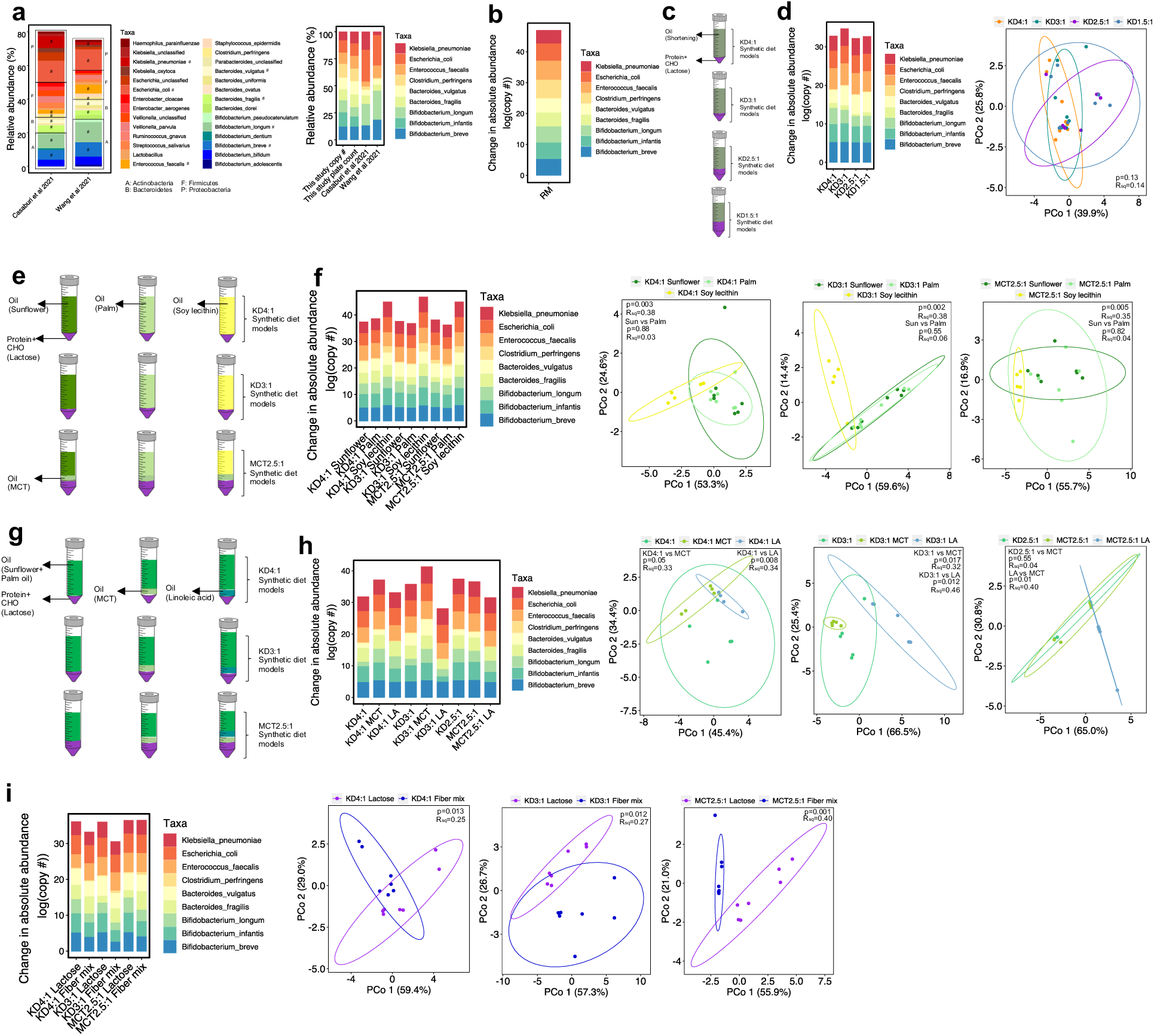
Effects of fat ratio, fat source/type, and carbohydrate source for KD-based synthetic culture media on metagenomic profiles of a model human infant microbial community. a. The bacterial species in the published data of infant gut microbiome, representing more than 1% relative abundance (left, the rectangular boxes donate phyla and the # denotes the species that were chosen for this study based on the highest relative abundance in their respective phyla) and bacterial species comprising the model human infant microbial community, as compared to published data from human infants (right) b. Change in bacterial species abundance after 24 hour culture in rich complex medium as a control (average of n=10) c. Experimental design: KD-based synthetic culture media was formulated with differing fat ratios for anaerobic culture of a model human infant gut microbial community d. Change in bacterial species abundance (left) and PCoA analysis of microbial taxonomic data (right) after 24 hour culture of model human infant gut microbial community in KD-based media with differing fat ratios (PERMANOVA, n=8/condition) e. Experimental design: KD-based synthetic culture media was formulated with differing fat sources that vary in level of saturation for anaerobic culture of a model human infant gut microbial community f. Change in bacterial species abundance (left) and PCoA analysis of microbial taxonomic data (right) after 24 hour culture of model human infant gut microbial community in KD-based media with differing fat sources (PERMANOVA, n=5-7/condition). g. Experimental design: KD-based synthetic culture media was formulated with differing fat types for anaerobic culture of a model human infant gut microbial community h. Change in bacterial species abundance (left) and PCoA analysis of microbial taxonomic data (right) after 24 hour culture of model human infant gut microbial community in KD-based media with differing fat types (PERMANOVA, n=5/condition). i. Change in bacterial species abundance (left) and PCoA analysis of microbial taxonomic data (right) after 24 hour culture of model human infant gut microbial community in KD-based media with differing carbohydrate sources (PERMANOVA, n=7/condition).

**Supplementary Figure 4.**
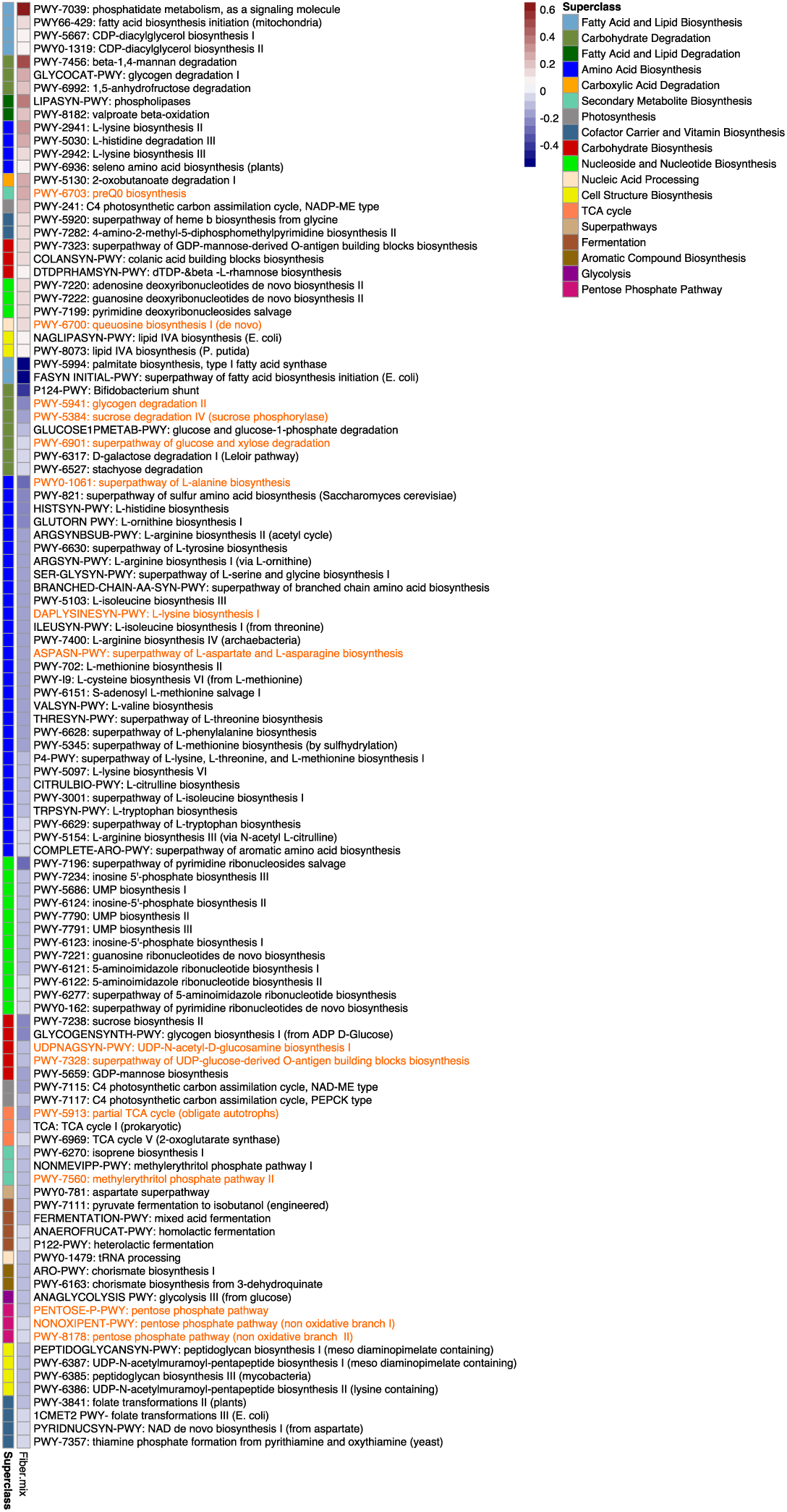
Addition of dietary fiber to KD-based synthetic culture media alters metagenomic signatures in a model human infant gut microbial community. Heatmap of differential metagenomic pathways (q<0.05) seen in model human infant gut microbial community after 24 anaerobic culture in fiber-containing KD-based media compared to lactose-containing KD-based media across all KD conditions (KD4:1, KD3:1, and MCT2.5:1, General Linear Model, n=21/condition).

**Supplementary Figure 5.**
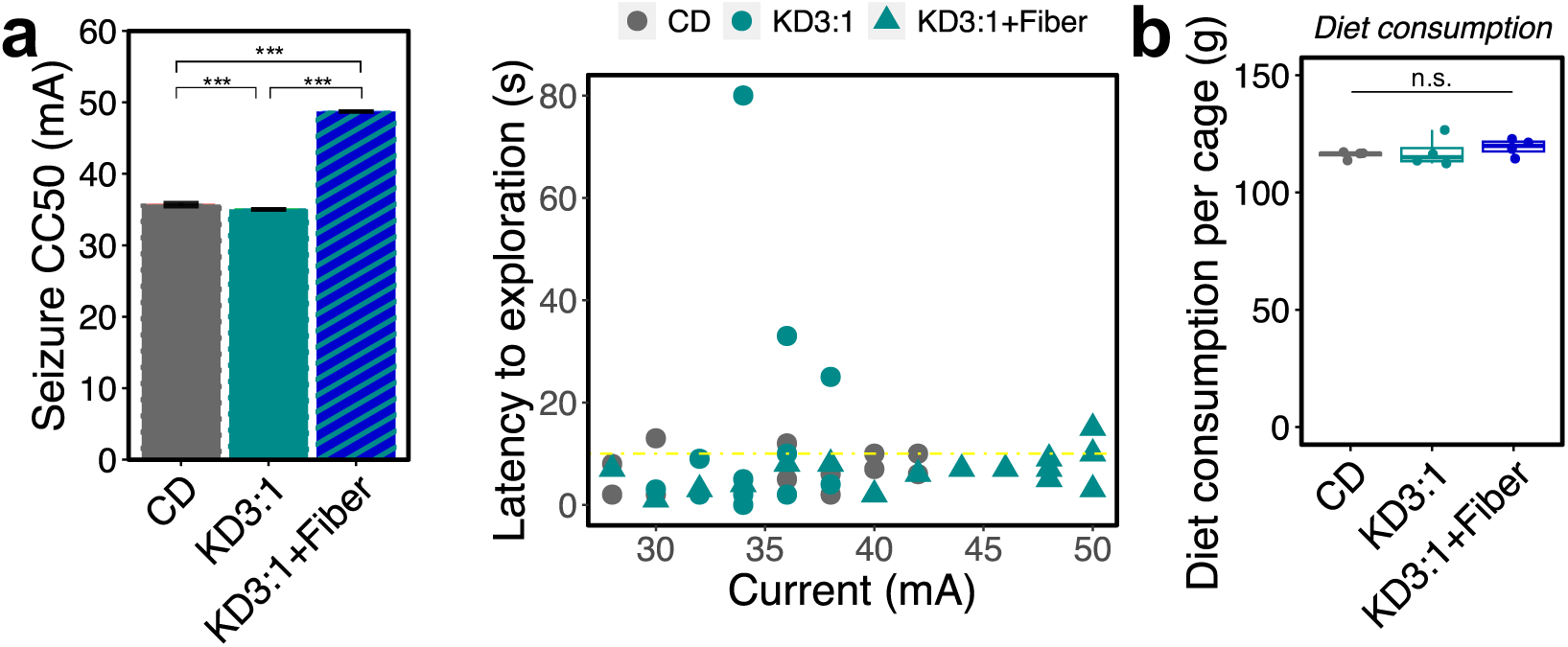
Addition of fiber to KD3:1 as a solid diet phenocopies increases in seizure resistance seen with liquid diet. a. 6-Hz seizure threshold (left) and latency to exploration (right) for mice fed KD3:1+fiber mix, KD3:1, or CD as solid diet (left, one-way ANOVA with Bonferroni, n=16 mice/condition, ***p<0.001). Yellow line at y = 10 s represents threshold for scoring seizures. b. Average consumption of solid diets (n=4 cages/condition; Kruskal-Wallis with Dunn’s test. Data are presented as box-and-whisker plots with median and first and third quartiles).

**Supplementary Figure 6.**
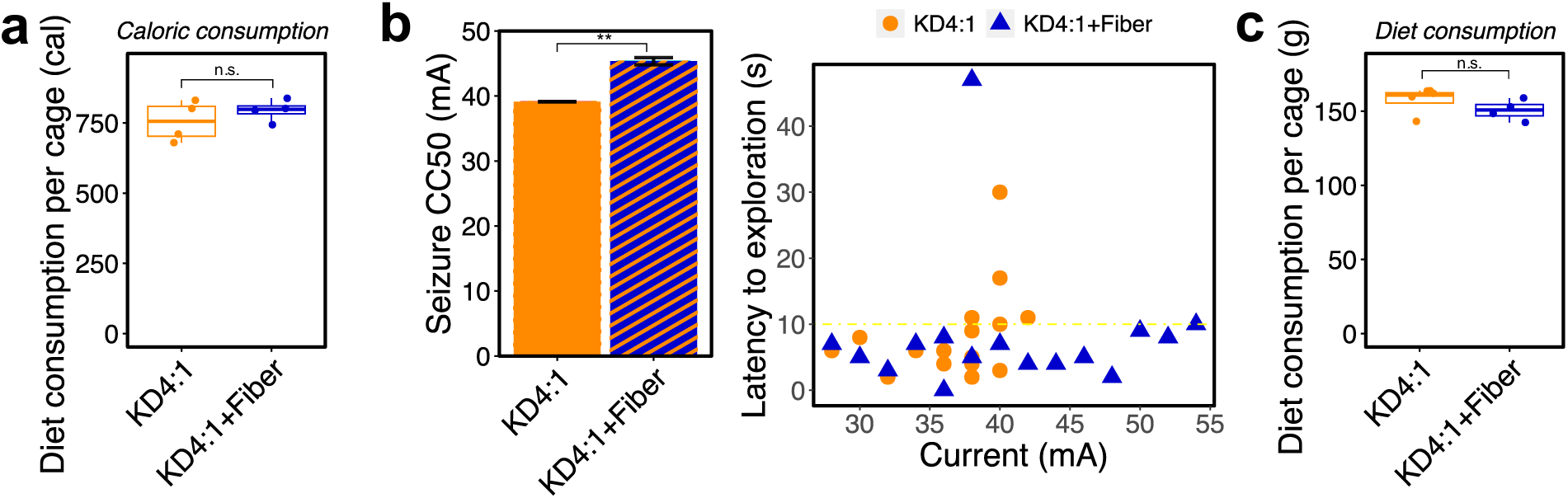
Addition of excess fiber to KD4:1 as a solid diet phenocopies increases in seizure resistance seen with liquid diet. a. Average caloric intake per cage for KD4:1 and KD4:1+fiber administered as liquid diet (n=4 cages; Wilcoxon signed-rank test. Data are presented as box-and-whisker plots with median and first and third quartiles). b. 6-Hz seizure threshold (left) and latency to exploration (right) for mice fed KD4:1+fiber mix or KD4:1 as solid diet (left, Welch’s t-test n=16 mice/group, ***p<0.001). Yellow line at y = 10 s represents threshold for scoring seizures. c. Average consumption of solid diets (n=4 cages; Wilcoxon signed-rank test. Data are presented as box-and-whisker plots with median and first and third quartiles).

**Supplementary Figure 7.**
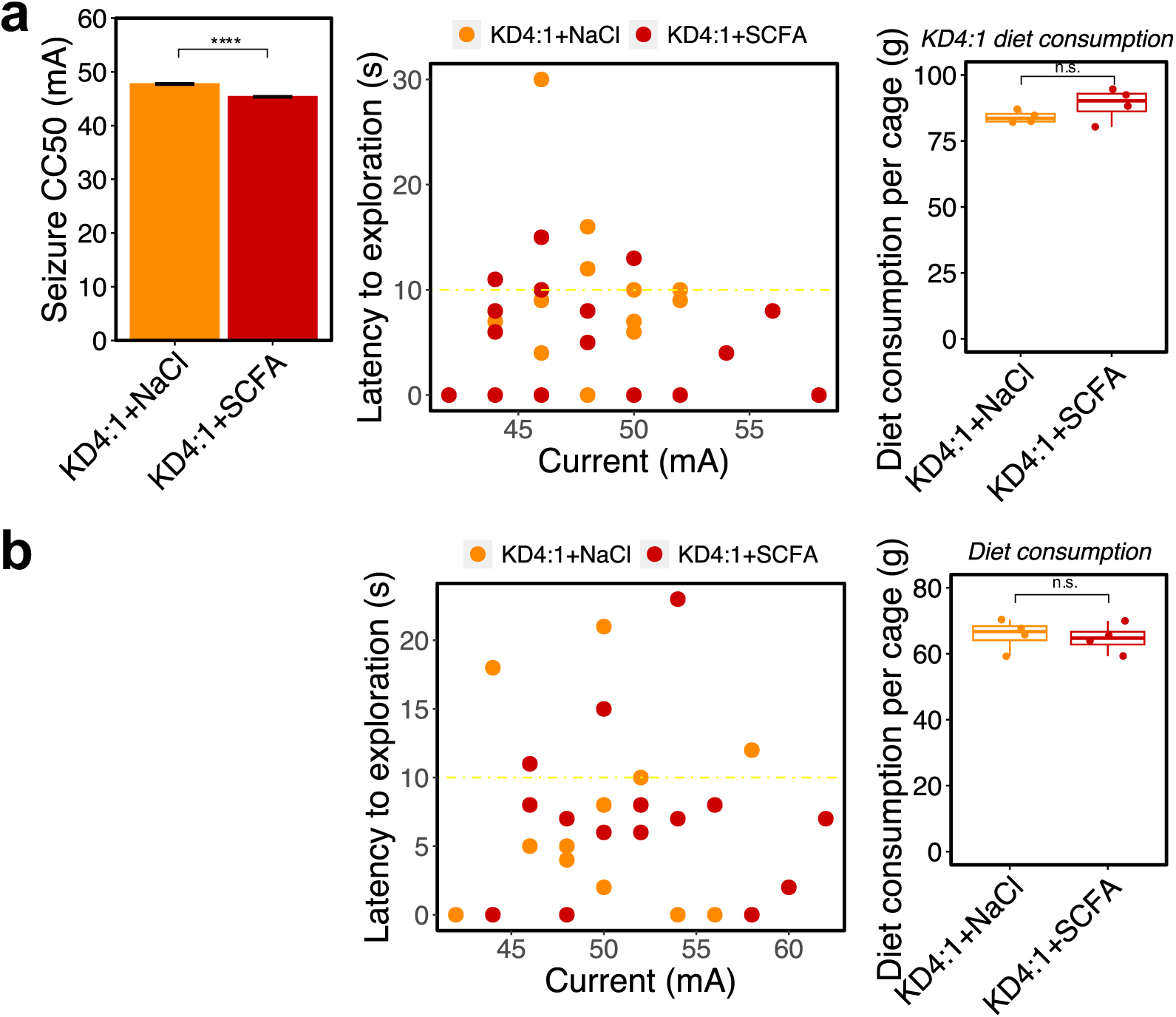
SCFA supplementation does not phenocopy effects of fiber supplementation on KD-induced response to 6-Hz seizures. a. 6-Hz seizure threshold (left) and latency to exploration (middle) for mice fed KD4:1 paste diet and supplemented with SCFAs or vehicle (NaCl) control in the drinking water (left, Welch’s t-test n=16 mice/group, ****p<0.0001). Yellow line at y = 10 s represents threshold for scoring seizures. Average consumption of paste diets (right, n=4 cages; Wilcoxon signed-rank test. Data are presented as box-and-whisker plots with median and first and third quartiles). b. 6-Hz seizure threshold (left) and latency to exploration (middle) for mice fed KD4:1 + SCFAs or vehicle (NaCl) control as a paste diet (left, Welch’s t-test n=16 mice/group, ***p<0.001). Yellow line at y = 10 s represents threshold for scoring seizures. Average consumption of paste diets (right, n=4 cages; Wilcoxon signed-rank test. Data are presented as box-and-whisker plots with median and first and third quartiles).

**Supplementary Figure 8.**
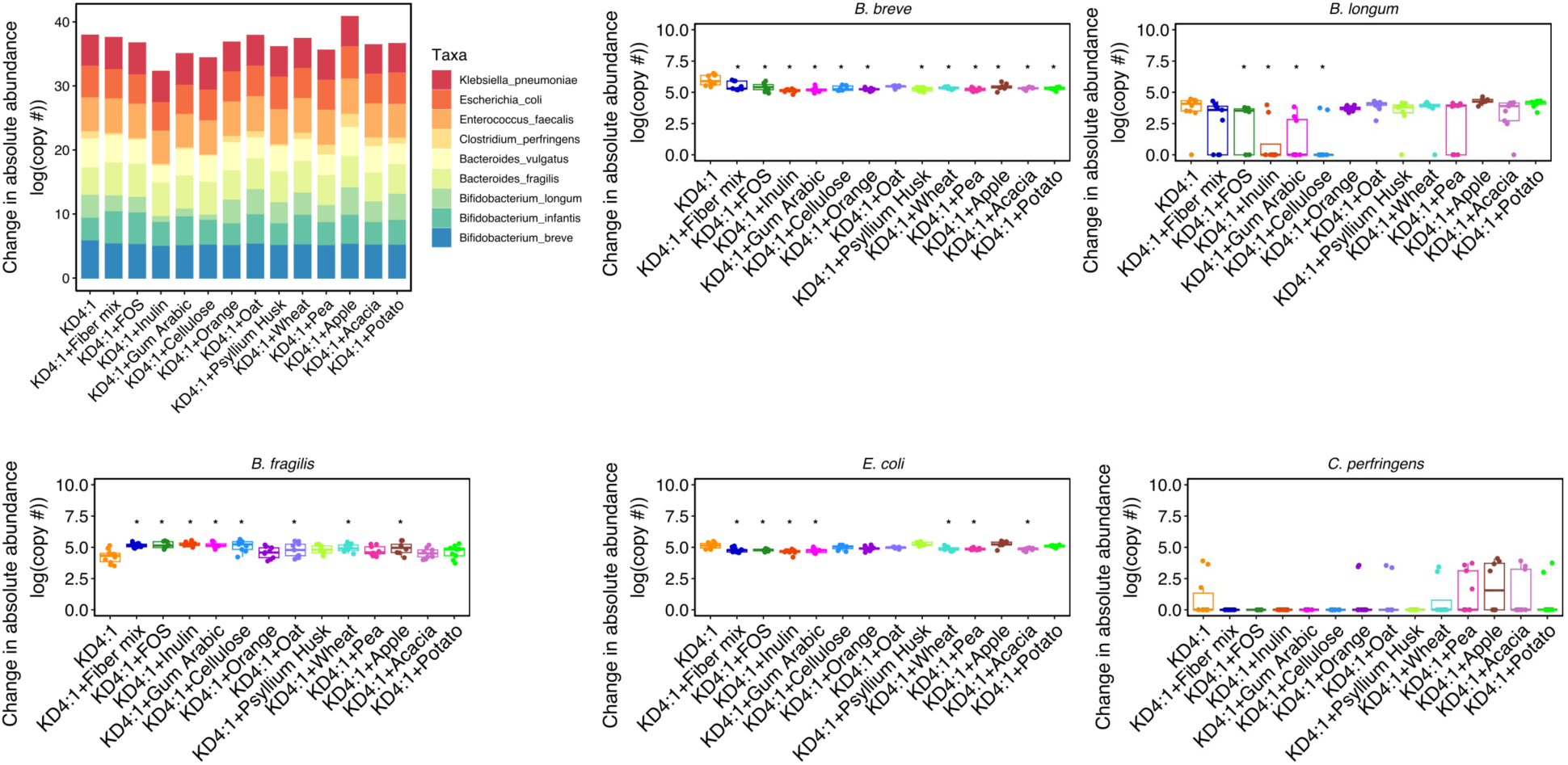
Supplementation of 13 dietary fiber sources and types to KD4:1 infant formula differentially alters the taxonomic composition of a model human infant gut microbial community. Change in bacterial species abundance after 24 hour culture of model human infant gut microbial community in KD4:1 infant formula with differing fiber sources and types, relative to KD4:1 alone (n=8-10. Kruskal-Wallis with Dunn’s test, ^∗^p < 0.05. Data are presented as box-and-whisker plots with median and first and third quartiles).

